# LiquidCNA: tracking subclonal evolution from longitudinal liquid biopsies using somatic copy number alterations

**DOI:** 10.1101/2021.01.05.425414

**Authors:** Eszter Lakatos, Helen Hockings, Maximilian Mossner, Weini Huang, Michelle Lockley, Trevor A. Graham

## Abstract

Cell-free DNA (cfDNA) measured via liquid biopsies provides a way for minimally-invasive monitoring of tumour evolutionary dynamics during therapy. Here we present liquidCNA, a method to track subclonal evolution from longitudinally collected cfDNA samples based on somatic copy number alterations (SCNAs). LiquidCNA utilises SCNA profiles derived through cost-effective low-pass whole genome sequencing to automatically and simultaneously genotype and quantify the size of the dominant subclone without requiring prior knowledge of the genetic identity of the emerging clone. We demonstrate the accuracy of liquidCNA in synthetically generated sample sets and *in vitro* and *in silico* mixtures of cancer cell lines. Application *in vivo* in patients with metastatic lung cancer reveals the progressive emergence of a novel tumour sub-population. LiquidCNA is straightforward to use, computationally inexpensive and enables continuous monitoring of subclonal evolution to understand and control therapy-induced resistance.

## Introduction

Liquid biopsies, primarily the analysis of cell free DNA (cfDNA) present in blood samples, offer the potential for regular longitudinal and minimally invasive monitoring of cancer dynamics [1, 2, 3, 4, 5, 6, 7]. Circulating cfDNA is released into the blood via apoptosis or necrosis of cells. Tumour-derived cfDNA in the blood is detectable from tumours as small as 50 million cells [8], it shows correlation with disease stage [9, 10], and offers the same diagnostic potential as tissue-based biopsies [7]. cfDNA is an aggregate of DNA shed from multiple locations and multiple malignant cells across the body and hence a single sample can provide a comprehensive overview of systemic disease. Consequently, cfDNA is an exceptional resource for non-invasive tracking of tumour composition and for monitoring response to therapy or clinical relapse.

Typically, cfDNA analysis has focused on the detection of driver gene single nucleotide variants (SNVs), with the size of mutation-bearing clones inferred from the relative sequencing read count at the mutation site. For instance, in high-grade serous ovarian cancer (HGSOC) the frequency of *TP53* mutation in cfDNA is a measure of tumour burden and is predictive of treatment response [11]. In colorectal cancer, *KRAS* mutation frequency in cfDNA is predictive of response to anti-EGFR therapy [4].

Somatic copy number alterations (SCNAs) are widespread in cancers [12, 13, 14], and have been used extensively to track tumour composition and dynamics over time [15, 16, 17, 18]. SCNAs can be detected in cfDNA without prior knowledge of the tumour SCNA profile, through measurement of the relative number of reads mapping within ‘bins’ spaced across the genome [19]. Relative differences in read count between bins can be sensitively detected even when the total read count is low [20, 21, 22], meaning that SCNAs can be detected with a fraction of the sequencing depth required for SNV detection. Therefore SCNA profiling offers a high-throughput and cost-effective means to evaluate cfDNA samples [23, 24, 25, 26, 27, 28].

Whilst measuring clone sizes based on the frequency of SNVs is straightforward, deriving quantitative information on the proportion of tumour population that carries a particular SCNA is challenging. Tumour cells are not the only contributors to the cfDNA pool, and an SCNA can in theory change the copy number to any non-negative integer value. Thus total read count per bin is a noisy compound function of the relative tumour cell contribution to the total cfDNA pool, and the specific copy number of the alteration.

Here we present a new method to identify and track tumour subclonal evolution based solely on measurement of SCNAs from longitudinal cfDNA samples. Our algorithm, named **liquidCNA**, firstly determines the contribution of tumour DNA to the total cfDNA pool (i.e. cellularity/purity) and then uses SCNA data to characterise and quantify the size of the most pervasive (putatively resistant) subclone emerging or contracting over time. The efficacy of the method is demonstrated using synthetic datasets, *in vitro* cell line mixtures, and *in vivo* via longitudinal analysis of cfDNA from lung cancer patients undergoing targeted treatment.

## Results

### Emergent subclone tracking from copy number information

First, we derive a mathematical definition of the problem of tracking an emergent (putatively resistant) tumour subclone from longitudinal cfDNA samples, typically taken throughout the course of treatment. We consider a tumour cell population undergoing continuous evolution characterised by two cell types, ancestral tumour cells (*A*) and an emerging subclone (*S*). We assume that liquid biopsies contain DNA originating from ancestral and subclonal tumour cells, as well as contaminating DNA from normal cells (*N*). The proportion of DNA arising from cells of the emergent subclone within the tumour is expressed by the subclonal-ratio, *r*_*i*_, while the overall proportion of tumour-originating DNA is termed the *purity* or tumour fraction of the sample, denoted by *p*_*i*_.

We consider that the copy number (CN) profile of each sample has been measured – for example using low-pass whole genome sequencing (lpWGS) – and so the genome can be divided into *segments*, contiguous regions of constant CN. Each measured segment CN in sample 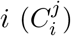 is the combination of each cell population’s CN at the *j*th genomic location (2 for normal cells and *C*(*A*) and *C*(*S*) for ancestral and subclonal tumour cells, respectively), weighted by the proportions of the three populations (Fig. 1).

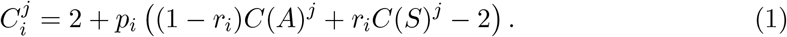

**Figure 1:**
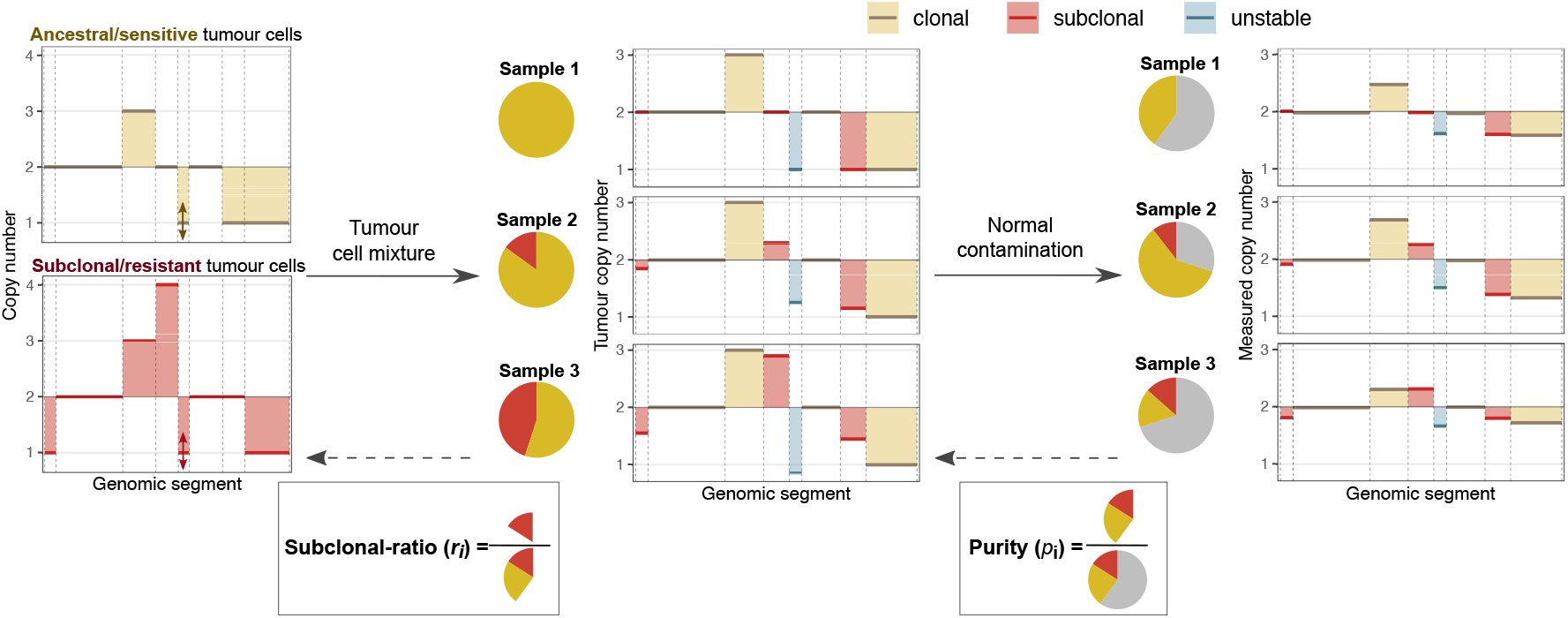
Schematic of copy number measurements. The first panel shows the SCNA profile of ancestral (in yellow) and subclonal (in red) tumour cells. At different sampling time-points, the overall tumour SCNA profile is a mixture of these profiles (second panel), influenced by the composition of tumour-derived DNA depicted on the pie-charts. Clonal, subclonal and unstable segments are indicated in yellow, red and blue, respectively. Note that the CN of clonal segments remains the same. In the liquid biopsies taken at each time-point, contamination from normal cells leads to ‘flattened’ measured SCNA profiles (last panel) due to normal cells having a neutral karyotype. This contamination affects the CN of each segment. Our aim is to estimate purity (*p*_*i*_) and subclonal-ratio (*r*_*i*_) based on clonal and subclonal SCNAs.

We assume that each segment can fall into one of three categories depending on its CN in ancestral and subclonal tumour cells. *Clonal* alterations (and unaltered segments) are at the same CN in both tumour populations, and their measured CN is only affected by the purity of a sample. *Subclonal* segments represent SCNAs that are unique to the emerging subclone. Their measured CN is influenced by the subclonal-ratio of a sample, as well as sample purity. Finally, segments that do not follow either of these patterns – due to uncertain measurements or ongoing instability – are termed *unstable*. Our aim is to estimate the underlying purity and subclonal-ratio, *p*_*i*_ and *r*_*i*_, from longitudinal CN measurements of clonal and subclonal segments (Fig. 1).

### Estimation of subclonal-ratio

Estimation is carried out in three steps (Fig. 2a and Methods). First, the *purity* of each sample is assessed using the distribution of segment CN values. We assume that the majority of segments have integer CN in all tumour cells, hence the distribution is expected to have distinct peaks at regular intervals of *p*_*i*_, corresponding to clonal segments with CN of 1, 2, 3, etc.(Fig. 2b). We derive the purity estimate as the value that minimises the squared error between observed and expected peaks (Fig. 2c). The inferred purity values are used to correct the segment CN values, thus estimating the tumour-specific CN of each segment.

**Figure 2:**
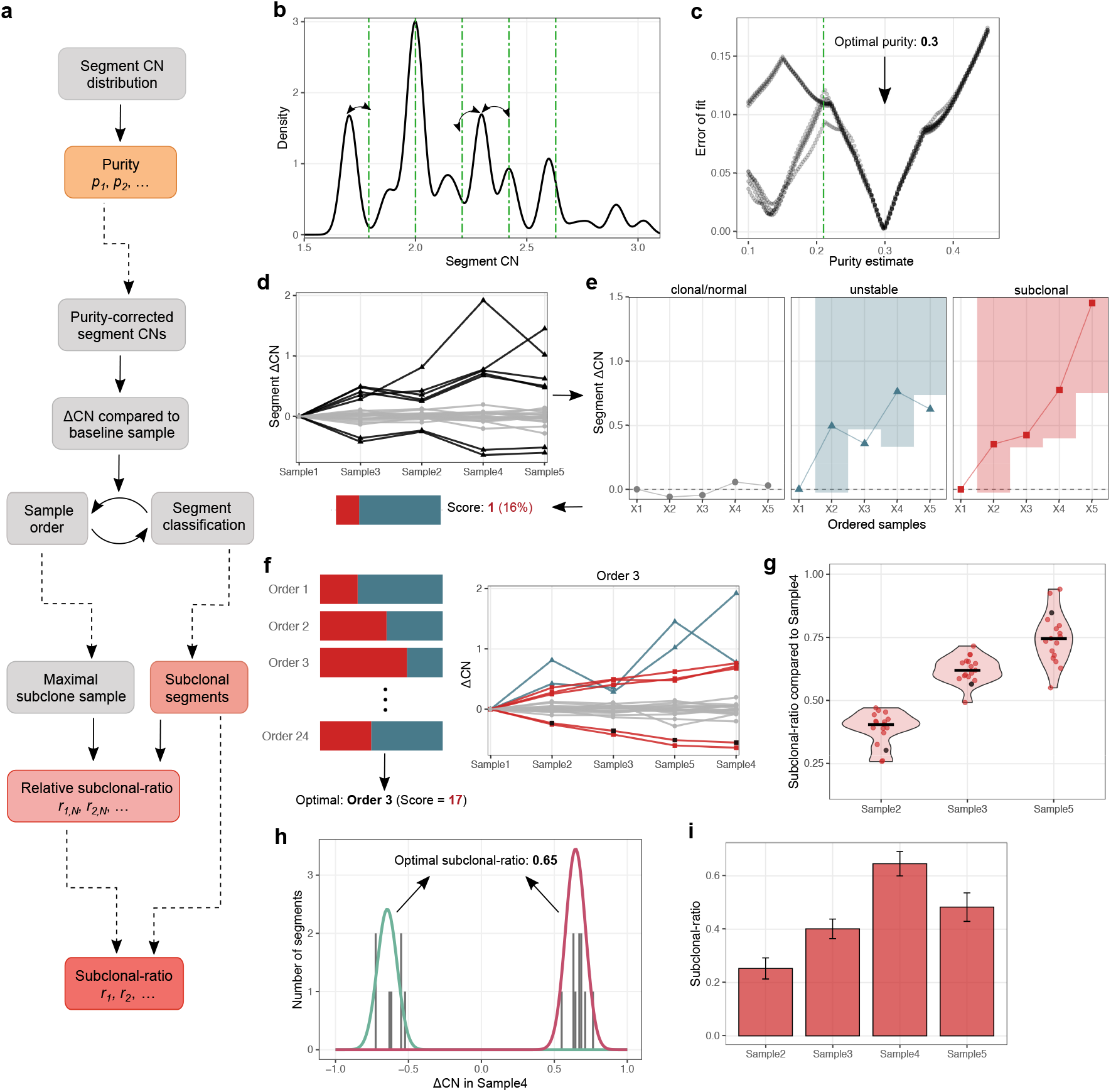
Illustration of the estimation algorithm. **(a)** Outline of the steps of the estimation algorithm. **(b)** Purity estimation based on the peaks of the distribution of segment CNs. Green lines show the peaks expected at an example purity of 0.21. **(c)** The error of a range of purity estimates, computed from the distance of observed and estimated peaks in (b). Each line corresponds to a smoothing kernel applied to the raw segment CN distribution. The optimal purity is indicated with arrow. **(d)** Change in segment CN values (ΔCNs) plotted according to an example sample order. The number of subclonal segments computed in (e) is indicated below. **(e)** Classification of segments based on the sample order in (d). Segments with low variance are classified as clonal (in grey). Non-clonal segments are evaluated whether they follow a quasi-monotone pattern (indicated by the shaded regions) and classified as unstable (outside of shaded region, in blue) or subclonal (in red). **(f)** ΔCN values plotted according to the optimal sample order maximising subclonal segments. Line colours indicate the class of each segment as in (e). **(g)** Relative subclonal-ratio estimation compared to maximal subclonal-ratio sample (right-most in (f)). Points show individual segment-wise estimates, with an example segment highlighted in black. Black line shows the median. **(h-i)** Subclonal-ratios and confidence intervals inferred by fitting a Gaussian mixture model to the ΔCN distribution of sub-clonal segments. The components of the best fit with means −*r* and *r* are shown in green and magenta in (h).

LiquidCNA does not require a mainly diploid tumour genome (i.e. major peak at CN = 2) to derive correct estimates, but will derive erroneous conclusions if the CN values – as measured by the CN quantification software, e.g. QDNAseq [19] – are incorrectly centred (e.g. major peak is defined as copy number 2, but the true value is copy number 3). To control for this an initial manual check of the CN profile is recommended prior to applying liquidCNA and renormalisation to the correct ploidy if required.

Next, for every segment we compute the change in CN, ΔCN, between each sample and a *baseline* sample that is assumed to have negligible proportions of the emerging (putatively resistant) subclone – for example a sample taken upon diagnosis or before start of therapy. ΔCN values naturally highlight subclone-associated segments altered in non-baseline samples, as these segments display markedly positive (CN gain compared to baseline) or negative (CN loss) values (Fig. 2d). From these ΔCNs we then establish the set of segments that are subclonal and the sample ordering that reflects increasing subclonal proportions. To do this, we examine each possible order of samples, classifying each segment as clonal (if the variance of its ΔCNs across samples is below a pre-defined threshold), subclonal (if it shows monotone change in ΔCN value along the order of the samples - i.e. if the ΔCNs are consistent with an emerging subclone) or unstable (if it does not correlate with sample order) according to that order (Fig. 2e). The order with the highest proportion of segments classified as subclonal is selected, and these subclonal segments are used for downstream computation of tumour composition (Fig. 2f). The methodology ensures that the *dominant* subclone associated with the most pervasive SC-NAs is evaluated and that subclonal-ratio inference is robust to segments with unstable CN.

Finally, we compute the relative and absolute subclonal-ratio of each sample using the identified set of subclonal segments. Relative subclonal-ratios are defined as the median ratio of segment ΔCNs compared to the sample with the maximum subclonal proportion (Fig. 2g). The absolute subclonal-ratio is computed based on the assumption that sub-clonal segment CN values correspond to distinct SCNAs that differ between ancestral and subclonal cells. The subclonal-ratio of sample *i* is therefore derived as the shared mean (*r*_*i*_) of a mixtures of Gaussian distributions with constrained means −*r*_*i*_, +*r*_*i*_, etc., fitting the ΔCN distribution of subclonal segments (Fig 2h). We also provide the 95% confidence interval of the absolute subclonal-ratio estimate based on the shared variance of the fitted Gaussians (Fig 2i).

LiquidCNA outputs both relative and absolute subclonal-ratio measures, since for most applications the relative value holds sufficient information on how the subclonal (putative resistant) population changes between time-points. Relative proportions are also less susceptible to the measurement noise in the measured segment CNs, while a combination of low subclonal proportion and high sequencing noise can cause the fitting of absolute subclonal-ratio estimates to fail to converge.

### Synthetic mixed populations

We first evaluated the performance of liquidCNA using *synthetic* datasets where input values of subclonal proportion and purity were known. We generated synthetic datasets characteristics matching typical longitudinal measurements of patients. In order to simulate imperfect measurements, we added varying levels of normally distributed measurement noise (defined by the dimensionless parameter *σ*) to bin-wise CN values (Fig. 3a-c and Methods).

**Figure 3:**
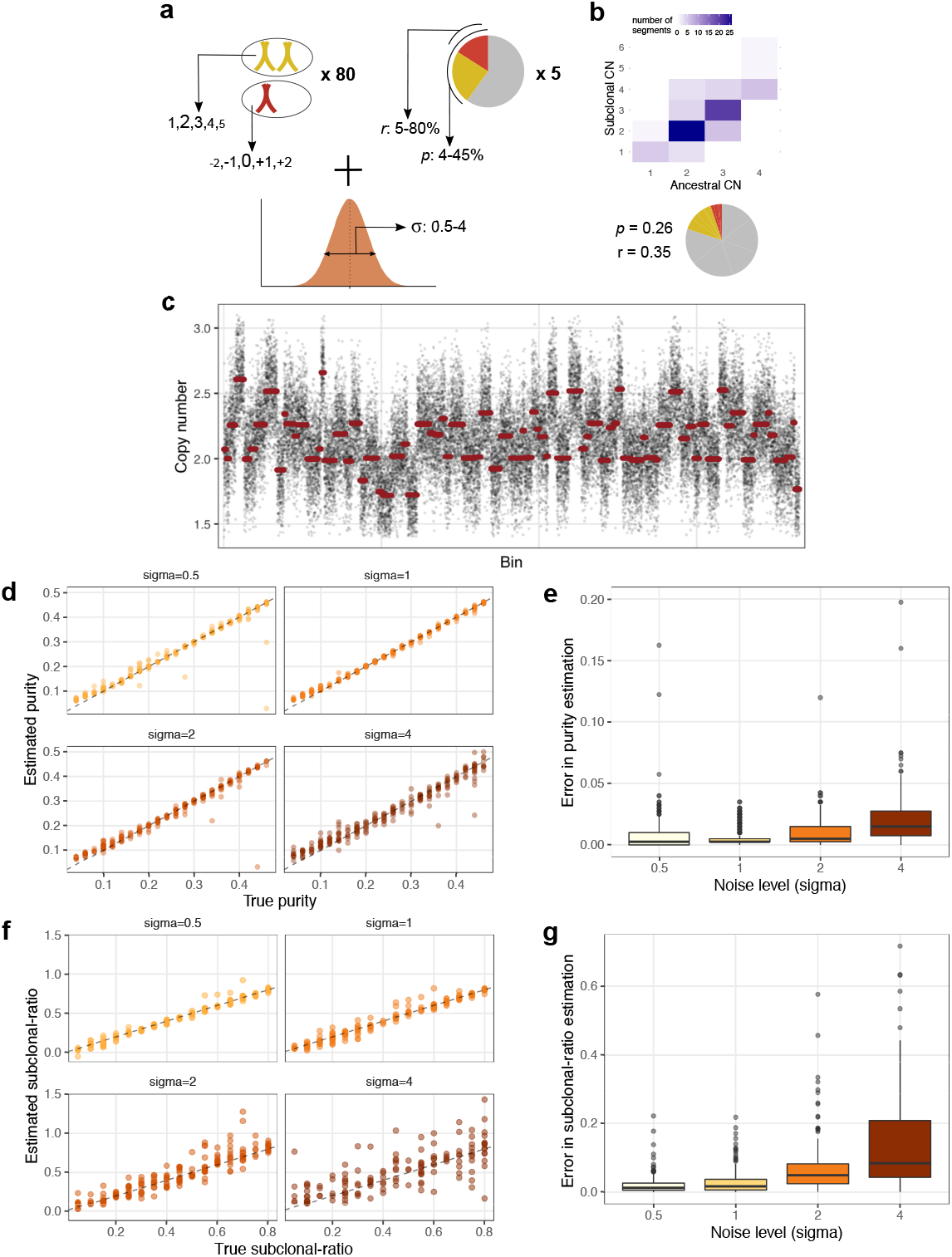
Estimation of mixtures of synthetic cell populations. **(a)** Parameters used to randomly sample synthetic datasets including simulated measurement noise. The font-size of copy number states indicates their probability. **(b)** A randomly generated sample. The heatmap depicts the distribution of segment CNs in ancestral and subclonal cells, and the proportion of cell populations is shown on the pie-chart (red: subclonal, yellow: ancestral, grey: normal). **(c)** Copy number profile of the sample in (b), with raw bin-wise and segmented copy number values shown in black and red, respectively. **(d)** Estimated purity of 1,000 synthetic samples with varying levels of noise (*σ*), plotted against the true theoretical purity. The *y* = *x* line is indicated with dashes. **(e)** Error of purity estimation (absolute difference to true purity) for samples with noise level indicated on the x axis. **(f)** True and estimated subclonal-ratio of 200 synthetic datasets (1,000 samples) with varying levels of noise (*σ*). **(g)** Error in subclonal-ratio estimation for datasets with increasing noise level. Box-plot elements in (e)(g) stand for: center line, median; box limits, upper and lower quartiles; whiskers, 1.5x interquartile range; points, outliers.

We evaluated the accuracy of the purity estimation on 250 synthetic samples (Fig. 3d), and found that purity p could be estimated within 2% of the true tumour fraction in 90% of samples at noise levels *σ* ≤ 1. The error on the purity estimation was greater when the noise was increased (Fig. 3e), and was most pronounced in samples with high noise and low tumour fraction. Consequently, we restricted our subsequent analysis to only cases of higher purity (*p*_*i*_ ≥ 0.1).

Next, we derived subclonal-ratios using purity-corrected CN profiles on the higher purity subset of synthetic mixtures. We set a threshold to filter out clonal segments (see Fig. 2e) such that at least 10 segments were retained and the proportion of retained segments classified as subclonal was maximal following segment classification. Fig. 3f shows the true and estimated subclonal-ratios for 50 synthetic experiments. Overall, we found that subclonal-ratio was estimated with ~ 5% error, and the accuracy was influenced by measurement noise (Fig. 3g). Relative subclonal-ratios (calculated compared to the sample with highest subclonal proportion) were estimated with higher accuracy (error ~ 3%, Fig. S1a-b). We found that computing absolute subclonal-ratios in a two-step process from these values yielded similar results to direct estimation by fitting a Gaussians mixture model, and provided an estimate even in cases where the direct estimation did not converge (Fig. S1c and Methods). The proportion of unstable segments, unlike noise, had little effect on the estimation accuracy (Fig. S2).

### Mixtures of ovarian cancer cell lines

Next, we evaluated liquidCNA on real data derived from *in vitro* mixtures of two paired high grade serous ovarian cancer (HGSOC) cell lines [29] (see Method and Table S1). HGSOC cells were ideally suited for this evaluation as high levels of chromosomal instability are a hallmark of the disease [30, 31]. We anticipated that liquidCNA will be most applicable for the tracking of subclonal evolution in malignancies with high CNA burden[26].

We divided a population of OVCAR4 cells into two aliquots, and the first aliquot was untreated and classified as ‘sensitive’. In a process described in detail by Hoare *et al.* [29], cells from the second aliquot were cultured so that they evolved resistance to platinum-containing chemotherapy and thus were termed ‘resistant’. In addition to the high SCNA burden inherited from the ancestral sensitive cell line, resistant cells acquired new SCNAs during the *in vitro* evolution of resistance (Figure 4a).

**Figure 4:**
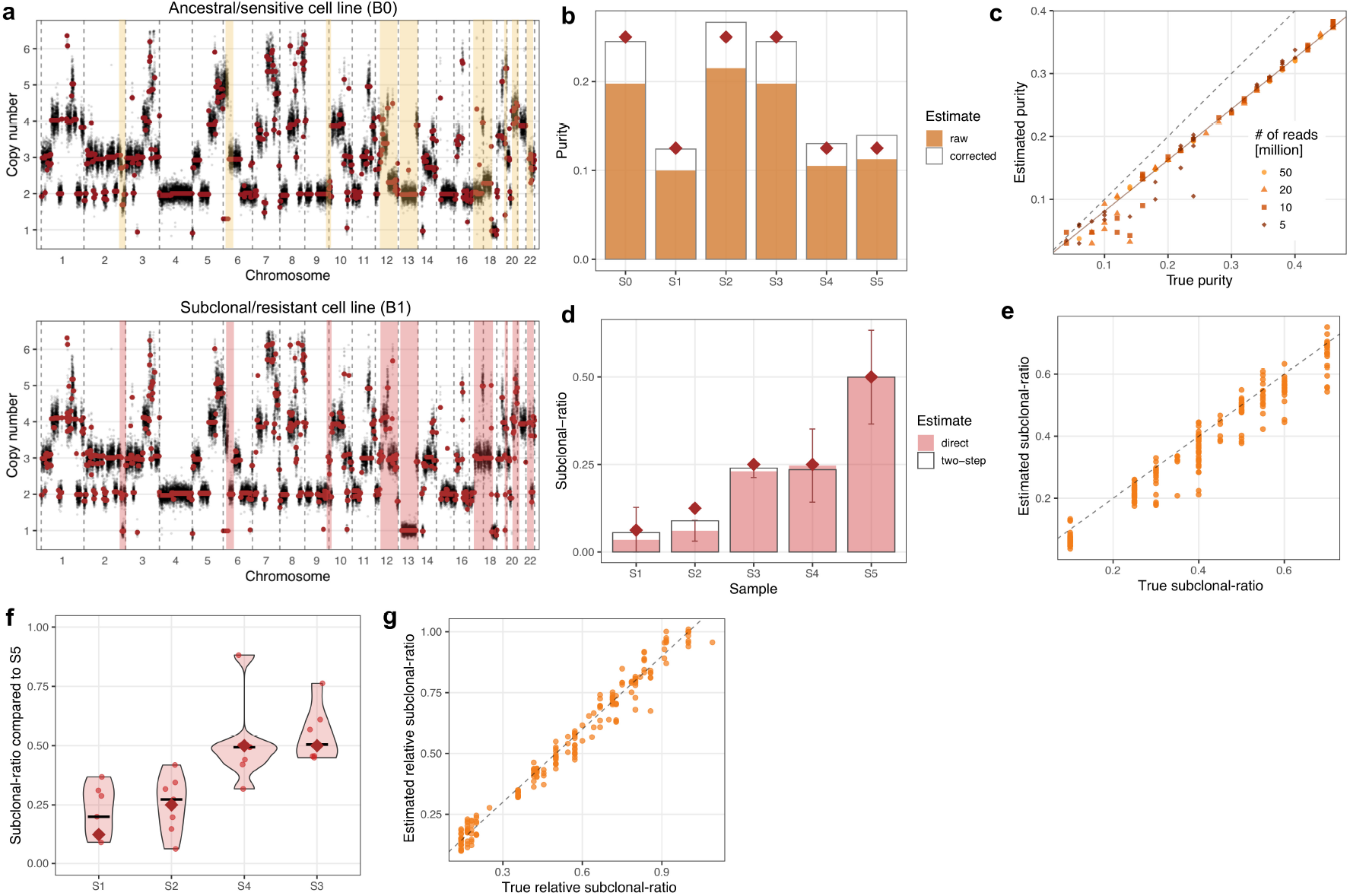
Estimation of mixtures of high grade serous ovarian cancer cell lines. **(a)** Copy number profile of the ancestral/sensitive and subclonal/resistant HGSOC cell lines. Raw bin-wise and segmented copy number values are shown in black and red, respectively. Resistant-specific subclonal SCNAs are highlighted. **(b)** Purity estimates of samples S0-S5. Corrected values are computed using the linear fit in (c). Theoretical purity values are indicated by maroon diamonds. **(c)** True (theoretical) and estimated tumour purity of 120 *in silico* HGSOC cell line mixtures. *y* = *x* and the linear fit of the estimates (*y* = 0.81*x*) are shown with dashed and solid lines, respectively. Point shape and shade indicate total number of reads per sample. **(d)** Subclonal-ratio estimates for samples S1-S5. Shaded and empty bars indicate estimates derived using direct (Gaussian fit) and two-step (from relative ratios in (f)) methods, respectively. Error bars show 95% confidence interval of the direct estimate, maroon diamonds indicate theoretical values. **(e)** True and estimated subclonal-ratio of 50 *in silico* datasets constructed of samples from (c) with 50 million reads. **(f)** Relative subclonal-ratio estimates for samples S1-S4, compared to S5. Estimates from each subclonal segment are shown with dots, the median estimates are indicated by black lines, and true values with maroon diamonds. **(g)** True and estimated relative subclonal-ratio in the 50 datasets shown in (g).

We then mixed, in varying known proportions, the genomic DNA extracted from the two cell lines, with sensitive cells representing the ancestral and resistant cells the emerging subclonal population. The mixtures were further diluted with DNA from blood samples of healthy volunteers assumed to have a diploid genome; this modelled the effect of normal contamination in patient samples (Table S1). These DNA mixtures were sequenced to mean depth 1.3x and composite SCNA profiles were generated (see Methods). In addition, we generated further *in silico* mixtures by sampling and mixing genome-aligned reads from sequencing data from each of the three cell types sequenced individually. In these mixtures, we controlled the total number of reads per sample to study the effect of variable read depth and associated measurement noise.

First, we used liquidCNA to estimate the purity of *in vitro* mixed samples (samples S0-S5). The purity of each sample was estimated to be lower than the theoretical mixing proportion (Fig. 4b). In the *in silico* mixed samples, we found that there was a strong linear relationship between estimated and true purity (Fig. 4c). The underestimation of purity in the samples might be explained by our definition of theoretical purity in the *in vitro* and *in silico* mixing procedure (respectively defined as proportion of DNA weight versus the proportion of read counts). A highly aneuploid genome will likely have a higher weight than a diploid genome, therefore mixing of equal weights results in a higher proportion of normal genomes than expected. Our purity estimates were in agreement with observed peaks of the CN distribution (Fig. S3a), further confirming that there was no bias in the estimation. By fitting a linear model to the estimates, the theoretical tumour fraction could be fully recovered, as illustrated by the ‘corrected’ estimates of samples S0-S5 (Fig. 4b). The number of reads (sequencing depth) did not systematically influence the accuracy of estimating tumour fraction, but purity estimates of samples with low tumour fraction were noisier at low read depth (Fig. 4c). In summary, liquidCNA provided an accurate estimate for purity values when true purity was above 10%. Decreased measurement accuracy below 10% purity is consistent with our observations on synthetic data and is similar to reported limitations of other methods quantifying tumour fraction from lpWGS cfDNA [24, 20, 22]. Therefore, for samples below 10% predicted purity, we advise to discard the sample from downstream analysis, although low-purity samples may be usable if a very accurate purity estimate can be derived by other means.

Next, we inferred the subclonal-ratio for cell line mixtures using purity-corrected ΔCN values, with sample S0 used as the baseline sample for both *in vitro* and *in silico* sample sets. We could correctly order cell line mixtures according to subclonal-ratios without any *a priori* information (Fig. S3b), and both absolute subclonal-ratio and relative subclonal changes were estimated on average within 2% and 3% of the true subclonal percentage (Fig. 4d,f). In particular, we noted that samples S4 and S3 were accurately estimated as having an equal subclonal-ratio, despite originating from different biological replicates with different tumour purity, which was reflected in the small confidence intervals of their estimates. We also note that even though there were no *truly unstable* segments in this dataset as measurements were not taken over time, three non-clonal segments were classified as such, probably due to higher noise in their measured CN value.

Using datasets of randomly selected *in silico* samples with 50 million reads, we confirmed that our algorithm could accurately infer the subclonal-ratio of samples, in particular when considering relative proportions (Fig. 4e,g). Although the estimation quality decreased with lower read counts (Fig. S4), in most cases the estimated absolute and relative subclonal-ratio was within 15% and 10% of the true subclonal proportion, respectively. Furthermore, we found that cases with high estimation error were typically caused by low-purity samples, which could be easily identified and removed without *a priori* information, as demonstrated in Fig. S5.

Using the known theoretical mixing values of tumour-DNA content – instead of data-derived estimates – to derive purity-corrected CN values increased the estimation error, especially in low read count samples (Fig. S6). This finding emphasises that non-diploid genomes might bias alternative measurement methods and internal consistency in the method of deriving sample characteristics (purity and subclonal-ratio) is crucial when assessing the dynamics of the subclonal population.

### Subclonal analysis of patient samples

We used liquidCNA to analyse emergent subclones in longitudinal cfDNA samples from patients with non-small cell lung cancer (NSCLC) undergoing therapy, as previously reported by Chen and colleagues [24]. The liquid biopsies were collected as part of the FIGARO study (GO27912, NCT01493843), a randomised phase II trial designed to evaluate the efficacy of pictilisib, a selective inhibitor of phosphatidylinositol 3 kinase [32]. Pictilisib or placebo was given in combination with standard chemotherapy regimen which was determined based on the subtype of NSCLC. Blood samples were taken at baseline (day 1 of the first treatment cycle) and at 6-week intervals up to the end of treatment (EOT). DNA was isolated from the plasma of liquid biopsies and sequenced using lpWGS to an average depth of 0.5x, as described in details in [24].

Chen *et al.* [24] identified several SCNAs in EOT samples that were absent at baseline and described several genes within these regions that might be associated with resistance. We sought to apply liquidCNA to these cases to corroborate their observations, and further to quantify the size of emergent subclones over time in these patients.

We obtained the lpWGS data (fastq files) and performed CN profiling (see Methods) on patients with cfDNA samples from ≥3 time-points (n = 32). We identified three patients (1306, 2760 and 3209) whose sample series fulfilled the following criteria: (i) had a cfDNA sample taken on the first day of therapy with purity above ~20%; (ii) and had at least two non-baseline samples with purity above ~20%. Patients 1306 and 3209 were in the experimental arm of the study, while patient 2760 was assigned to the control arm; and all three patients have progressed during the course of the trial.

We ran liquidCNA on data from the three selected patients (discarding samples with purity below 10% (Fig. S7)) and examined the genomic segments that liquidCNA identified as subclonal relative to baseline samples (Fig. 5).

**Figure 5:**
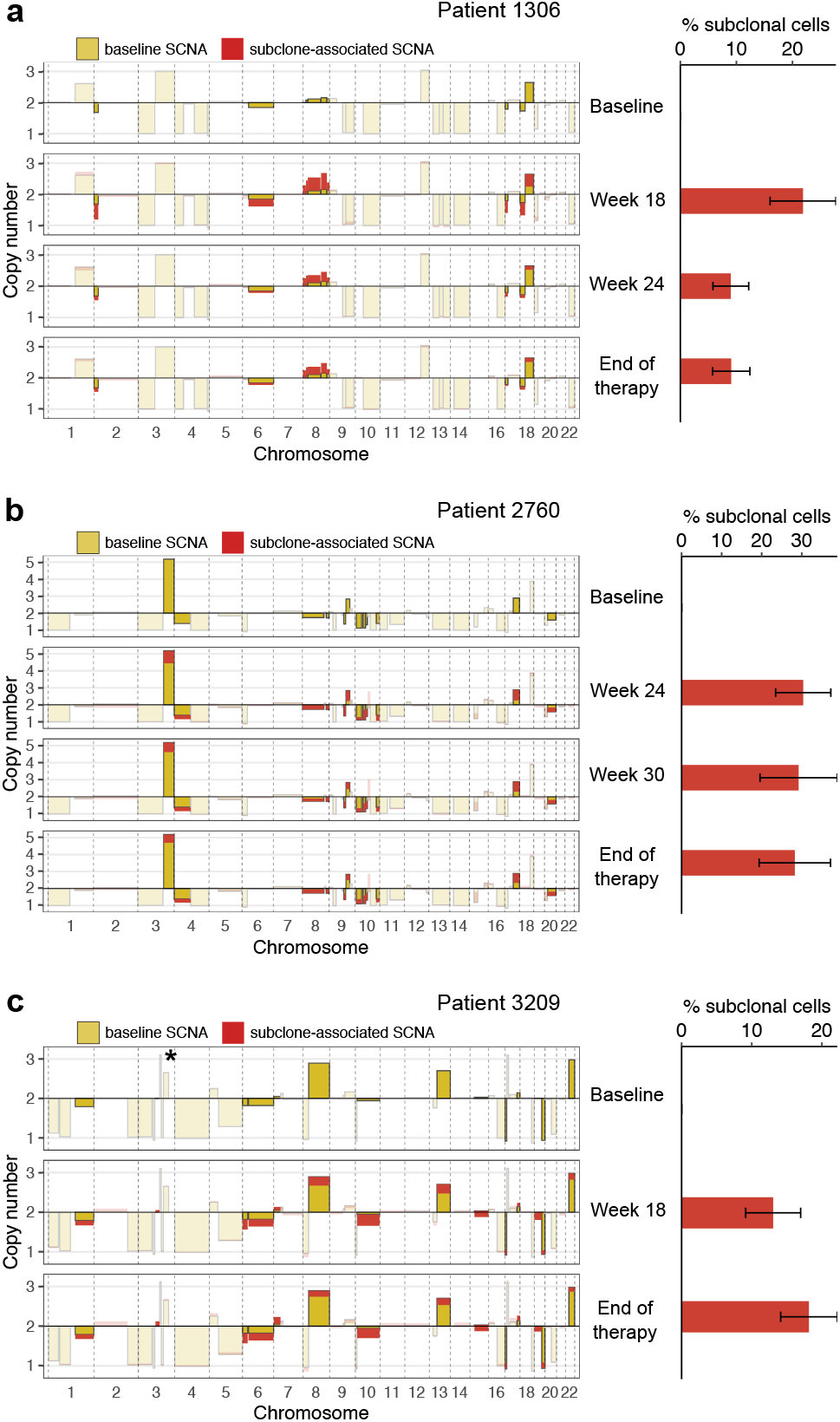
Estimation in cfDNA samples from patient data. Subclone-specific copy number changes and subclonal-ratio in lung cancer patients (**a**) 1306, (**b**) 2760, and (**c**) 3209 from [24]. Left: purity-corrected SCNA profiles. Yellow bars show the CN of each segment in the baseline sample, and red bars indicate subclonal deviations from this value in non-baseline samples. Regions of subclone-specific CNAs are also indicated by darker shades. Right: estimated resistant proportion of each sample with 95% confidence intervals. Note that only samples with >10% purity were analysed (c.f. S7). A bar of CN=6 on chromosome 3 (indicated by asterisk) has been omitted from (c) for better visualisation.

While we observed a good overlap with the CNs previously reported to be associated with subclonal evolution through therapy (Figures 5 and S8 of [24]), we also found a few segments that were missed or additionally identified by liquidCNA. The original study focused on the comparison of pre- and post-treatment and highlighted SCNAs occurring between the first and last time-points. As our analysis put equal focus on all time-points, it classified some of the previously identified segments as unstable if the CN progression was not consistently with subclone evolution. Furthermore, some segments were too small to pass our initial filtering. On the other hand, liquidCNA was able to identify subclonal segments which were at an abnormal CN in the baseline sample, and subsequently showed diploid CN or a further gain/loss in subclonal tumour cells. For example, in the samples from Patient 1306, whilst liquidCNA identified subclonal SCNAs on chromosomes 2, 6 and 8 that overlapped with the findings of the original study; it also detected additional subclonal changes on chromosomes 17 and 18. However, we did not observe the previously described focal loss on chromosome 7 (harbouring the gene *MLL3*), probably due to its small size. Overall, we identified 10, 12 and 17 subclone-associated SCNAs in patients 1306, 2760 and 3209, respectively. A further 6 segments in patient 2760 were classified as non-clonal but ‘unstable’ as the CN over time was not consistent with the pattern defined by the emerging subclone. As samples from patient 2760 had lower purity, these inconsistent CN changes might have resulted from measurement noise.

We found that the emerging subclone accounted for 10 to 30% of the tumour derived DNA in the cfDNA in the three patients evaluated. Patient 2760 showed evidence of a subclonal proportion consistently around 30%, which could be explained by samples from this patient taken at later time-points. Samples from patient 3209 obtained at 18 weeks and end of therapy contained below 20% DNA derived from subclonal tumour cells (Fig. 5). Patient 1306, on the other hand, showed a contracting subclone that reduced in proportion from 20% presence at week 18 to <10% at the end of therapy. In case the total population size was known – which might be accessible from additional measurements of the tumour-associated cfDNA pool –, the tumour subclone fractions established here could also be converted into growth rates to enable future predictions of the tumour dynamics.

## Discussion

We present liquidCNA, a computational algorithm to infer longitudinal subclonal dynamics using copy number measurements. Our algorithm performs simultaneous analysis of several longitudinal samples to identify sample purity, subclonal SCNAs and the abundance of an emerging subclone. LiquidCNA distinguishes between SCNAs that are associated with the emerging subclone and those showing unstable behaviour, and consequently is not confounded by uncertain CN measurements.

We validate our method both on synthetic SCNA datasets, and *in vitro* and *in silico* mixtures of two ovarian cancer cell lines. We successfully infer the proportion of the dominant subclone in all of the above datasets, with good accuracy across a range of sample qualities defined by the noise level or sequenced reads. In patients with lung cancer, liquidCNA applied to lpWGS data derived from longitudinal liquid biopsies (cfDNA) shows the emergence of subclones during therapy and identifies genomic regions associated with the emergent tumour cells.

We demonstrate that liquidCNA can identify and quantify emerging subclones from cfDNA samples, therefore enabling tracking of tumour subclone evolution through the course of therapy. Deciphering the evolutionary trajectory of cancer can aid prognostic and therapeutic decision-making and further our understanding of therapy-induced drug resistance [33]. Measuring the dynamics of tumour composition is particularly crucial for prospective monitoring during an adaptive therapy regime aiming to control resistant subclones [34, 35, 36]. Furthermore, the proportion of cfDNA that is tumour-derived (what we term ‘purity’) in itself is a promising biomarker for determining initial therapy response and prognosis [5, 37], as well as for tracking tumour progression during and after therapy [2, 10, 6, 24].

We note that there are limitations in our liquidCNA method. Since our inference relies on heterogeneous copy number profiles and subclone-specific SCNAs, we cannot analyse cancer (sub)types with very low chromosomal instability, for example microsatellite unstable tumours. Conversely, extremely high levels of ongoing instability might bias our analysis due to the lack of stable subclone-associated SCNA profile, and therefore liquidCNA is not suitable for oligo-metastatic disease if spatially separate metastases carry distinct karyotypes. Furthermore, the accuracy of our estimation reduces at low purity (below 10%). However, a tumour fractions above this regime were observed in a substantial number of patients, especially in late stage disease where liquidCNA can offer the largest benefit, [24, 20, 6, 9, 22, 38]. In addition, recent studies have shown that the unique fragment length of tumour-derived cfDNA can be utilised to enrich for tumour purity either experimentally or bioinformatically [39, 40, 41]. Finally, liquidCNA tracks a single dominant subclone associated with the largest set of subclone-specific SCNAs, and if there are multiple smaller subclones (with less or no associated SCNAs), these will be ignored by the algorithm.

In summary, we provide a robust tool to derive quantitative information about dynamic changes in clonal composition from SCNA measurements derived from cfDNA. LiquidCNA enables real-time non-invasive tracking of subclonal tumour evolution, which can provide new insights into the evolution of SCNAs and the dynamical emergence of therapy-associated resistance.

## Acknowledgements

We thank Ann-Marie Baker for reviewing the clarity of the text, and Steve Gendreau and Craig Cummings from Genentech, Inc. for providing access to patient cfDNA sequencing results and for their critical comments on the presentation of the data.

This work was supported by the Wellcome Trust (grant 202778/Z/16/Z to T.A.G.) and Cancer Research UK (grant A19771 to T.A.G. supporting E.L.; Advanced Clinician Scientist Fellowship C41405/A19694 to M.L.; Clinical Research Training Fellowship to H.H.). M.L. also received support from a Barts and The London Charity Strategic Research Grant (467/2244). T.A.G. also received founding from the National Institutes of Health, National Cancer Institute (grant U54 CA217376).

## Author contributions

E.L., W.H., M.L. and T.A.G. conceived and designed the study. M.L. and T.A.G. acquired funding for the study. E.L. developed the inference method and performed bioinformatic analysis. H.H. and M.M. performed *in vivo* experiments and sequencing. E.L. and T.A.G. wrote the original draft, and all authors reviewed and approved the manuscript.

## Competing interests

The authors declare no competing interest.

## Methods

### Formal definition of the problem

#### Copy number measurements

We consider a tumour that consists of two distinct cell populations, ancestral (*A*) and subclonal (*S*) tumour cells, and continuously sheds cell-free DNA (cfDNA) into the blood circulation. A typical scenario would be ancestral cells representing drug-sensitive tumour cells present before cancer therapy, and subclonal cells denoting the emerging subclone with resistance to therapy. The proportion of DNA originating from these two cell types changes over time as we take measurements via blood samples (Fig. 1). Since cell-free DNA found in blood can also originate from normal (non-tumour) cells of the body, the measured DNA is contributed by a mixture of the two tumour cell populations (*A* and *S*) and normal cells (*N*). At each time-point *i* the proportion of these three populations in the measured sample, *s*_*i*_, depends on the proportion of all tumour-derived DNA (the *purity* of the sample, *p*_*i*_) and the proportion of subclone-derived DNA from the tumour (the subclonal-ratio, *r*_*i*_):

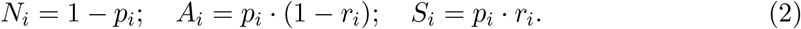

Our aim is to track the dynamics of the subclonal (putatively resistant) population by determining the subclonal-ratio for each time-point, *r*_*i*_, or the change in subclonal-ratio between time-points, *r*_*i*_/*r*_*k*_ = *r*_*ik*_. To this end, we use the copy number values as typically measured by lpWGS of the sequential cfDNA samples.

Let us consider distinct genomic regions with homogeneous copy number state, *segments*. We assume that the copy number (CN) state of most segments stays constant over time in a particular population. Therefore the *j*th segment is characterised by a set of three time-independent absolute CN states, *C*(*N*)^*j*^, *C*(*A*)^*j*^, *C*(*S*)^*j*^, corresponding to the local CN in normal, ancestral and subclonal cells, respectively. The copy number of segment *j* as measured in the *ith* sample, 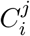 is the combination of these three absolute CNs, weighted by the proportions of DNA derived from the three cell populations at that time-point (*N*_*i*_, *A*_*i*_, *S*_*i*_). We know that normal cells are at a diploid state, hence *C*(*N*)^*j*^ = 2 for all *j*. Therefore, using the purity and subclonal-ratio defined in (Eq. 2),

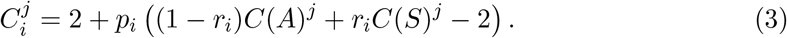

Since all cells in a cell population share the absolute CN for a given segment, the values *C*(*S*)^*j*^ and *C*(*A*)^*j*^ are always integers. Therefore in theory, measured CNs from a given sample should be limited to a discrete set of values defined by these integer states, making it possible to solve the set of equations formed by (Eq. 3) for *p*_*i*_ and *r*_*i*_ using linear algebra.

However, we have to take into account that all real sequencing measurements have a level of imprecision introducing variation on top of this relationship. Using the term *σ*_*ij*_ to represent the noise in the *i*th measurement of segment *j*, (Eq. 3) becomes,

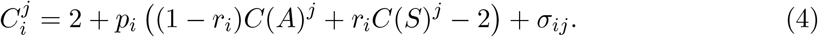

with the magnitude and family of this noise depending on the specifics of the technology used for CN measurement, especially the sequencing depth [19]. This measurement noise – associated with a continuous distribution – broadens the set of 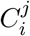 values, rendering a linear algebra solution impossible. Hence, our aim becomes to derive an inference of *p*_*i*_ and *r*_*i*_ despite this unknown noise, *σ*_*ij*_.

#### Segment classification

Each segment can fall into three categories depending on their respective copy number states in the two types of cells. (i) *Clonal* segments have the same absolute CN in ancestral and subclonal tumour cells, *C*(*A*)^*j*^ = *C*(*S*)^*j*^. A special case of clonal segments are segments of *neutral* CN, where *C*(*A*)^*j*^ = *C*(*S*)^*j*^ = 2. (ii) *Subclonal* segments have different absolute CNs in the ancestral and subclonal tumour population, *C*(*A*)^*j*^ ≠ *C*(*S*)^*j*^. These segments represent SCNAs that distinguish the *subclone* from its ancestor, even though they are not necessarily associated with a selective/phenotypic difference (e.g. drug-resistance) directly. (iii) *Unstable* segments are neither clonal nor associated with the emergent subclone, and therefore are best described by a time-dependent tumour-wide CN value, 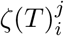 that does *not* depend on *r*_*i*_. These segments can arise if a genomic region cannot be measured reliably or if on-going genomic instability introduces novel SCNAs during the time tracked by our samples. We can assume that the number of such segments is small compared to the total number of measured segments.

Depending on whether segments are clonal, subclonal or unstable, their measured CN across samples will change according to the subclonal-ratio and purity of each sample. For simplicity, we omit the term *σ*_*ij*_ and its derivatives, but the reader should keep in mind that all equations are subject to measurement noise:

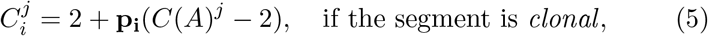

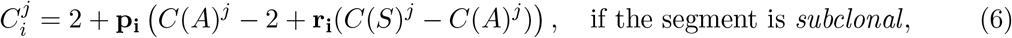

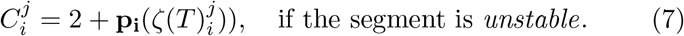

Figure 1 illustrates how the measured CN of segments depend on the parameters *r*_*i*_ and *p*_*i*_ highlighted above. In the following sections, we use (Eqs. 5) & (6) to estimate the underlying parameters, *p*_*i*_ and *r*_*i*_, via three steps (Fig. 2).

### Estimation algorithm

#### Purity estimation

Purity estimation is carried out based on clonal (including neutral) segments. In general, we expect the majority of segments to fall into this category. Consequently, for the majority of segments their measured copy number follows (Eq. 5). Since *C*(*A*)^*j*^ can take only integer values, the distribution of segment CNs is expected to have distinct peaks at regular intervals of *p*_*i*_.

Using a peak-finder algorithm on the smoothed distribution of measured CN values, we directly compare the peaks to the values expected at a given purity, {2 − *p*_*i*_, 2, 2 + *p*_*i*_, 2 + 2*p*_*i*_,…}, as shown in Fig. 2b. The error of the fit to a purity, *p*_*i*_, is evaluated as the summed squared distance between each peak and the closest observed peak,

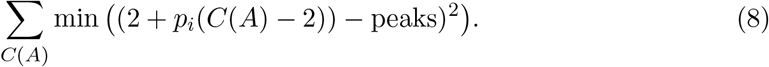

As the detected peaks of the data depend on the smoothing kernel used on the distribution, we perform this computation for a wide range of smoothing bandwidths (0.5×−2.5× the default value) and derive the purity estimate, 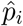, as the value that minimises the mean and/or median error across the range (Fig. 2c).

Then, we use the derived 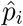 to re-normalise the measured copy number values and thus eliminate normal contamination. We gain an estimate of the tumour-specific CN 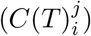 a mixture of ancestral and subclonal CNs:

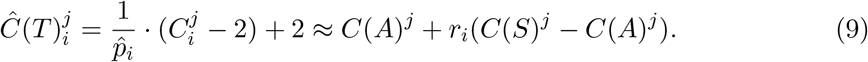

Note that, due to the noise in measurements, peaks from close absolute CNs can become indistinguishable in low-purity samples. Therefore we expect purity values below 5% to be indistinguishable (unless high sequencing depth is available) and also advise to discard samples with low purity (typically *p*_*i*_ < 0.1) as erroneous purity estimations can bias downstream computation.

#### Identifying subclonal segments and sample order

Next, we aim to identify the subset of segments with subclone-specific *subclonal* SCNAs that reflect the changes in subclonal-ratio over time. To easily assess the *change* in segment CNs, we designate a sample as *baseline*, and compute the change in segment CN, ΔCN, between each sample and this baseline sample. Typically, the sample taken upon diagnosis or before start of therapy (usually the first time-point, *s*_1_) can be used. We can assume that this sample has no or only negligible population of the emerging subclone, and therefore represents a pure ancestral population:

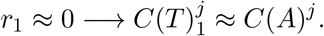

Hence the change in CN of a subclonal segment compared to the baseline becomes,

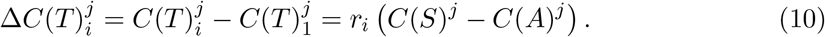

Furthermore, (Eq. 10) provides an informative quantity even if the baseline sample is not pure, as 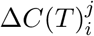 nonetheless describes the *change* in subclone-specific SCNAs.

In order to uncover which segments are truly *subclonal*, and how the subclonal-ratio changes over measurements, we need to identify a pervasive pattern across samples, and the subset of segments that consistently follows it. If the samples were taken so that the subclonal population increases over time-points, this pattern would be a monotone increase or decrease for all segments with subclone-specific SCNAs. While we cannot assume that the samples are taken in order of increasing subclonal proportions (e.g. a change of treatment between sampling times might lead to fluctuating population size in a resistance-associated subclone), we can aim to re-arrange them to follow this rule.

Consequently, we rephrase our aim as deriving (i) a *set of subclonal segments* that follow a monotone pattern across ordered samples; and (ii) an *ordering of samples* that is correlated with by the maximum number of (subclonal) segments. Formally, we are looking for a subset of segments, {*j*_1_, *j*_2_,…} and a permutation of samples (starting from the designated baseline sample), *s*_1_, *s*_*i*_,…, *s*_*N*_, where for every segment *j* ∈ {*j*_1_, *j*_2_,…}

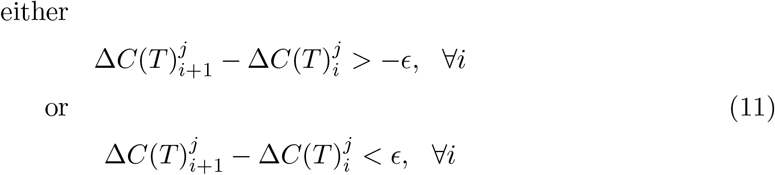

holds for all *i* for a pre-defined accuracy level, *ϵ*. We use an *ϵ* > 0 accuracy level to allow for samples with near-equal subclonal-ratio measured with uncertainty. We find that, for typical lpWGS datasets, *ϵ* ≈ 0.02–0.05 works well to account for the underlying measurement noise.

Figs. 2d-f illustrate the derivation of optimal sample order and subclonal segment set. We first separate clonal segments: since these have relative CN values of 0, apart from some measurement noise, we filter out any segment that has a standard deviation below a pre-defined threshold. We then evaluate (Eq. 11) over all remaining segments and over all orderings of the samples. As we expect 4-6 time-points per dataset, an exhaustive search of all possible permutations is feasible. Given a permutation, each segment is classified according to whether it follows (Eq. 11) – these are candidate subclone-specific and unstable segments, respectively (Fig. 2e). The optimal sample order is defined as the permutation that maximises the number of subclonal segments (Fig. 2f).

#### Subclonal-ratio estimation

Finally, we use the set of segments identified as *subclonal*, and compute the subclonal-ratio of each time point. We derive the (absolute) subclonal-ratio, *r*_*i*_, for each sample using (Eq. 10). As both *C*(*A*)^*j*^ and *C*(*S*)^*j*^ are assumed to be integers, and we know that *C*(*A*)^*j*^ ≠ *C*(*S*)^*j*^,

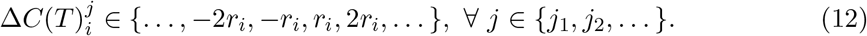

To take into account that the measured ΔCNs compared to the baseline, 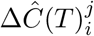 are influenced by noise, we fit these values with a mixture of Gaussian distributions where the mean of the Gaussians follows (Eq. 12), as illustrated in Fig 2h. The subclonal-ratio of a sample is derived as the constrained mean parameter, *r*_*i*_, of the Gaussian mixture optimising the fit (Fig. 2i). The 95% confidence interval of the inferred subclonal-ratio is computed based on the (shared) variance of the fitted constrained Gaussians.

The measurement noise propagated from segment CNs can lead to high spread in values, making estimates less robust and rendering the resolution of low subclonal-ratios (*r*_*i*_ ≤ 0.1) challenging, occasionally leading to the Gaussian-fitting step to fail. Therefore we also derive *relative* subclonal-ratios, which allow for a more general application not limited to good quality samples. In particular, relative values are compared to the maximal sample since its subclonal-ratio is assumed to be the most robust against measurement noise. We compute the relative deviation of each normalised subclonal tumour segment CN,

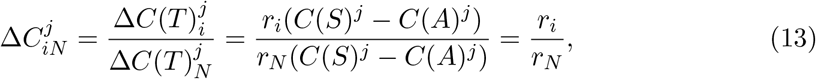

giving rise to a distribution of relative subclonal-ratio estimates (Fig. 2g). We derive a point estimate for the relative *r*_*i*_ of each sample as the median of this set,

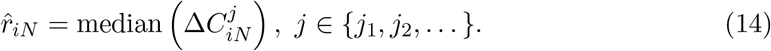

Absolute subclonal-ratio estimates can then be derived using these relative estimates in a two-step estimation process (as opposed to the direct estimation above): we derive *r*_*N*_ based on (Eq. 12), and subsequently compute *r*_*iN*_ · *r*_*N*_ to retrieve *r*_*i*_.

#### Generating synthetic datasets

We constructed synthetic datasets of 80 segments (of length varying between 120 and 800 bins) and 5 time-points as illustrated in Fig. 3a. For each segment, we generated sensitive segment copy number states (*C*(*S*)^*j*^) by randomly sampling from {1, 2, 3, 4, 5}, with neutral and close-to-neutral states occurring with higher frequency. Subclone-specific absolute CNs (*C*(*S*)^*j*^) were assigned by randomly sampling from *C*(*A*)^*j*^ + {−2, −1, 0, 1, 2}, with no change (giving rise to clonal segments) having a higher weight. For each sample, *s*_*i*_, we assigned purity and subclonal-ratio randomly from the ranges 0.04 < *p*_*i*_ < 0.45 and 0.05 < *r*_*i*_ < 0.8, with the exception of the baseline samples, where *r*_1_ < 0.04. We then recreated the measurement procedure of computing noise-ridden raw CN values in a given segment, *j*, by adding a normally distributed noise. The magnitude (standard deviation) of the noise was controlled by the noise level parameter, *σ* (representing differences arising from e.g. sequencing depth) and the CN of the segment (reflecting higher variance in higher CN states):

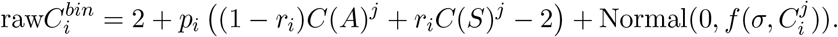

The final CN value of each segments, 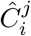 was computed as the mean of all 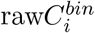 contained in the segment. In addition, we selected 2.5-15% of segments as *unstable*, and re-sampled their tumour-specific CN value to be independent of *r*_*i*_. Fig. 3b-c show parameters of a synthetic sample and its copy number profile.

#### Generating *in vitro* and *in silico* cell line mixtures

HGSOC cell line OVCAR4 was obtained from Prof Fran Balkwill (Barts Cancer Institute, UK) and grown in DMEM media containing 10% FBS and 1% penicillin/streptomycin. A resistant/subclonal HGSOC cell line (Ov4Cis) was generated by culturing an aliquot of the ancestral OVCAR4 cell line in increasing concentrations of cisplatin. For further details on cell culture and the celll lines, see [29].

We then extracted genomic DNA from both cell lines and from blood samples from healthy volunteers using QIAamp DNA Micro Kit (Qiagen, Hilden, Germany). Genomic DNA from the three sources was mixed in varying proportions (Table S1), measured as the mass of DNA inputted from each source, to a total of 20 ng DNA per sample and subjected to sonication with the Covaris M220 system. Libraries were prepared using the NEBNext Ultra II kit (New England Biolabs, Hitchin, United Kingdom) with 4 cycles of PCR amplification, indexed with unique dual indexing primers and sequenced on Illumina NovaSeq 6000 to a mean depth of 1.3x.

*In silico* mixtures were generated by bioinformatically mixing sequencing reads of DNA derived from the ancestral/sensitive, subclonal/resistant tumour cell lines and healthy blood cells. Similarly to synthetic samples, for each *in silico* sample we randomly assigned purity, 0.1 < *p*_*i*_ < 0.45, and subclonal-ratio, 0.05 < *r*_*i*_ < 0.8. We then sampled reads (using `samtools view −s`) from aligned read (bam) files of ‘pure’ ancestral, subclonal and normal samples (B0, B1 and N0) in proportions to match *p*_*i*_(1 − *r*_*i*_), *p*_*i*_*r*_*i*_ and 1 − *p*_*i*_, respectively. We also varied the total number of reads per sample (as a proxy for sequencing depth and consequently measurement noise), and generated 30-30 samples with 50, 20, 10, and 5 million total reads each.

#### Processing lpWGS samples

Fastq files derived from lpWGS samples (generated via sequencing cell line mixtures or obtained from [24]) were aligned to the human reference genome (version *hg19*, using bwa). We then processed bam files using the QDNAseq R package [19] employing DNAcopy for segmentation [42]. QDNAseq produced two copy number values for each genomic bin: a *raw* pre-segmentation and a *segmented* value grouping bins of equal CN together. The CN of bins on the pre-defined blacklist of QDNAseq and of those with <75% mappability was set to NA. Raw and segmented CN values for all cell line samples are available from https://github.com/elakatos/liquidCNA_data

Since QDNAseq returns normalised CN values (with neutral state at 1), we multiplied all values by 2 before proceeding with the estimation algorithm and re-normalised segment CN values to be centred at 2 exactly. We then re-defined segment boundaries using the ensemble of samples as regions of constant CN in *all* samples. This way break-points present in only a sub-set of samples (such as a subclone-specific SCNA) gave rise to segments handled separately for all samples. Updated segments with length below 6 mega-bases (120 bins of 50kb (cell line mixtures) or 12 bins of 500kb (patient cfDNA samples)) were excluded from the downstream analysis to filter out short segments sensitive to localised measurement biases.

Finally, we curated each segment CN by discarding bins with the most extreme 2.5% of raw segment values, and re-calculating the segment CN value as the mean of normal distribution fitted to the remaining raw CNs. We found that this curation had negligible effect for most segments, but successfully improved assigned segment CN values for more error-prone genomic regions.

## Data availability

Aligned sequencing data from HGSOC cell lines and in vitro mixtures (listed in table S1) are available from the European Nucleotide Archive (accession PRJEB42332). Raw and post-segmentation copy number values for these samples are available from https://github.com/elakatos/liquidCNA_data.

## Code availability

Estimation functions of liquidCNA implemented in R (version 4.0.3), an illustrative example in a Jupyter notebook and code generating and analysing synthetic and *in silico* data are available from https://github.com/elakatos/liquidCNA.

## Notes

### Competing Interest Statement

The authors have declared no competing interest.

### Summary of Updates

The author list and affiliations have been updated to reflect contributions from Weini Huang.

