## Supplementary Table and Figures for "LiquidCNA: tracking subclonal evolution from longitudinal liquid biopsies using somatic copy number alterations"

for

| Sample | Read count | Normal | Ancestral | Subclonal |
|---|---|---|---|---|
| B0 | 98 million | 0% | 100% | 0% |
| B1 | 76 million | 0% | 0% | 100% |
| N0 | 82 million | 100% | 0% | 0% |
| S0 | 87 million | 75% | 25% | 0% |
| S1 | 80 million | 87.5% | 11.72% | 0.78% |
| S2 | 70.5 million | 75% | 21.875% | 3.125% |
| S3 | 102 million | 75% | 18.75% | 6.25% |
| S4 | 83 million | 87.5% | 9.375% | 3.125% |
| S5 | 70.5 million | 87.5% | 6.25% | 6.25% |

Table S1: Proportion of DNA originating from normal, ancestral/sensitive and subclonal/resistant cells in the HGSOC cell line samples. Samples S0-S5 and *in silico* mixtures of B0, B1 and N0 are analysed in Figure 4.

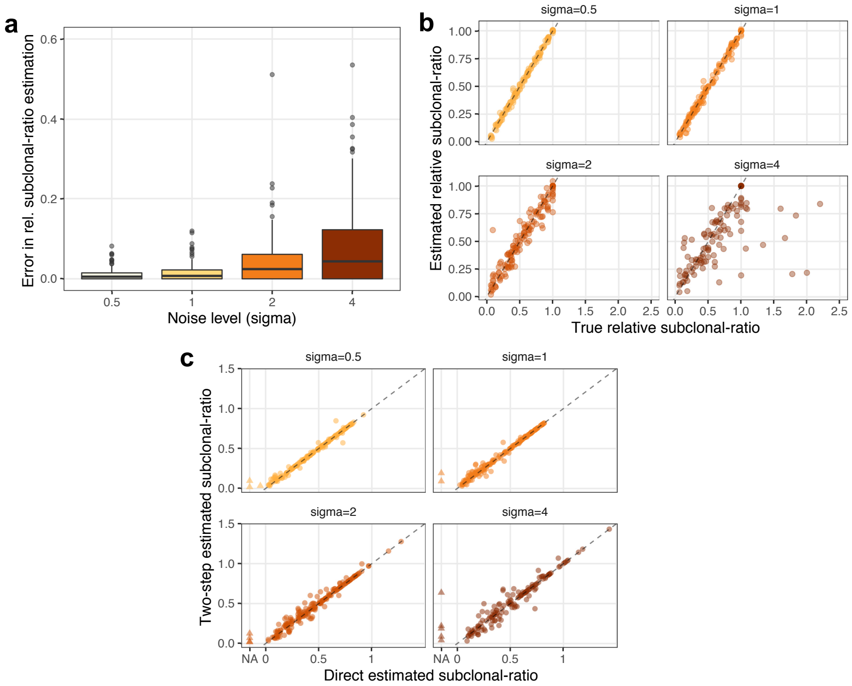


Figure S1: Subclonal-ratio estimation of synthetic mixtures using relative subclonal-ratios. **(a)** Error in relative subclonal-ratio estimation (compared to the sample with the maximal subclonal proportion) for 200 datasets with increasing noise level ($\sigma$), indicated on the x axis. Box-plot elements represent: center line, median; box limits, upper and lower quartiles; whiskers, 1.5x interquartile range; points, outliers. **(b)** True and estimated relative subclonal-ratio of the synthetic datasets in (a). **(c)** Subclonal-ratio estimates of the samples in (a) computed directly from Gaussian fits (x axis) or in a two-step estimation via relative subclonal-ratios (y axis), at different levels of $\sigma$. Samples where the direct estimation failed are indicated with triangles at value NA.

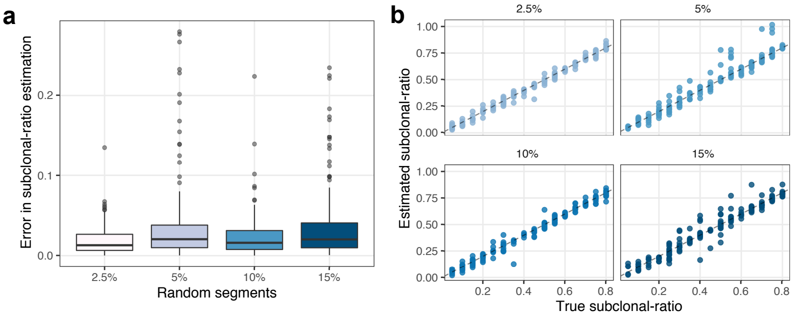


Figure S2: Subclonal-ratio estimation of synthetic mixtures with varying number of random segments. **(a)** Error in the subclonal-ratio estimate of 200 synthetic datasets with $\sigma = 1$ and the proportion of segments with random CN values indicated on the x axis. Box-plot elements represent: center line, median; box limits, upper and lower quartiles; whiskers, 1.5x interquartile range; points, outliers. **(b)** True and estimated subclonal-ratio of the synthetic datasets in (a).

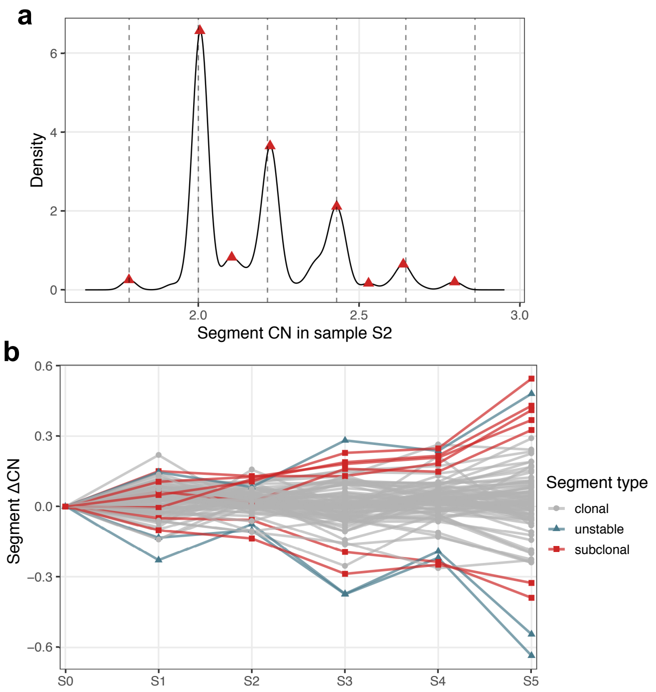


Figure S3: Estimation details of HGSOC cell line mixtures. **(a)** The distribution of measured segment CN values in sample S2. Peaks used in purity estimation are highlighted by red dots. Dashed vertical lines indicate the expected peak locations at the best fitting purity estimate, $\hat{p}_{S2} = 0.215$. **(b)** $\Delta$CN values of samples S0-S5, compared to the baseline sample, S0. Samples are shown in the order associated with increasing subclonal proportion, and coloured according to segment class (grey: clonal, blue: unstable, red: subclonal).

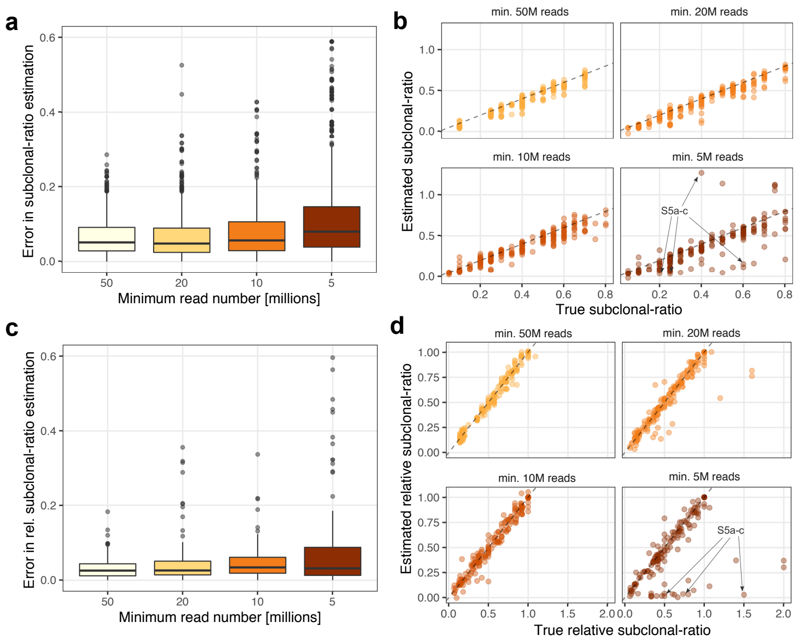


Figure S4: Subclonal-ratio estimation of *in silico* cell line mixtures with varying read counts. **(a,c)** Error in absolute (a) and relative (c) subclonal-ratio estimation for 200 datasets constructed of *in silico* samples with minimum required read count indicated on the x axis. Box-plots represent: center line, median; box limits, upper and lower quartiles; whiskers, 1.5x interquartile range; points, outliers. **(b,d)** True and estimated absolute (b) and relative (d) subclonal-ratio of the synthetic datasets in (a) and (c). The dataset indicated by arrows is re-analysed in Figure S5.

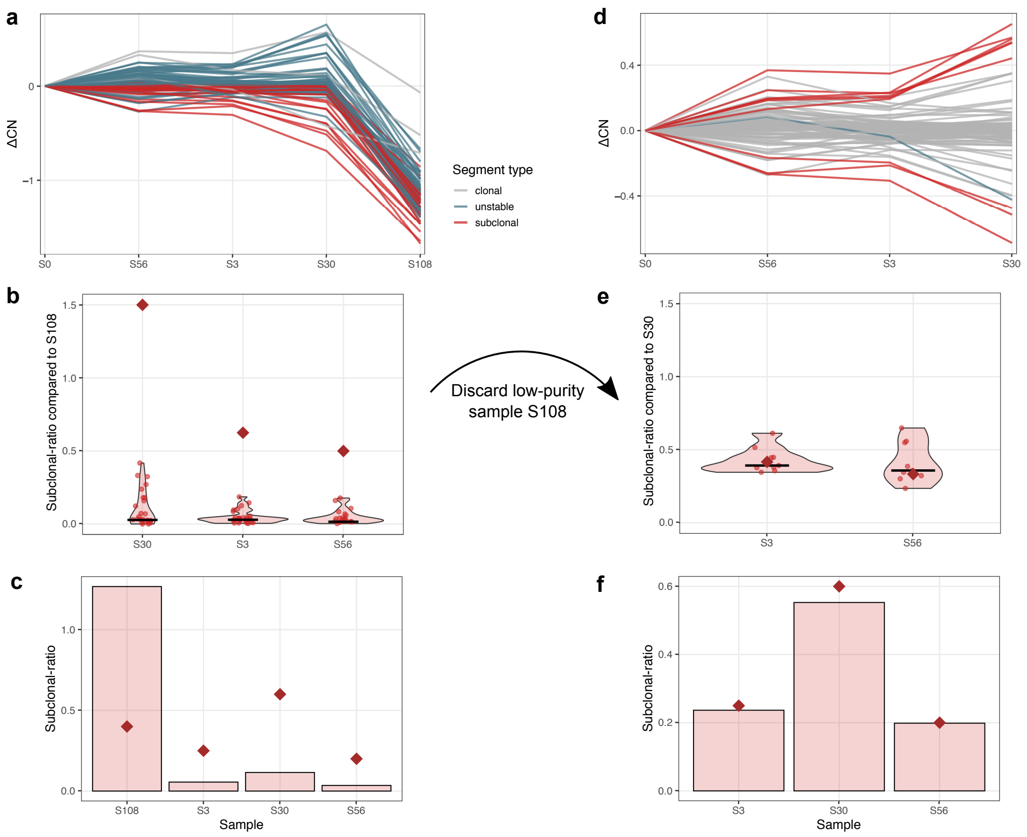


Figure S5: Manual curation of in silico cell line mixture dataset. **(a,d)** ΔCN values and classification with automatically assigned ordering of samples (a). As sample S108 clearly shows incorrect patterns due to low purity, it is discarded from downstream analysis (d). **(b,e)** Relative subclonal-ratio distribution computed for the original (b) and curated (e) maximal sample. Estimates from each segment are shown with dots, the median estimates are indicated by black lines, the true relative ratios with maroon diamonds. **(c,f)** Absolute subclonal-ratio estimates of the original (c) and curated (f) datasets. True values are indicated with maroon diamonds.

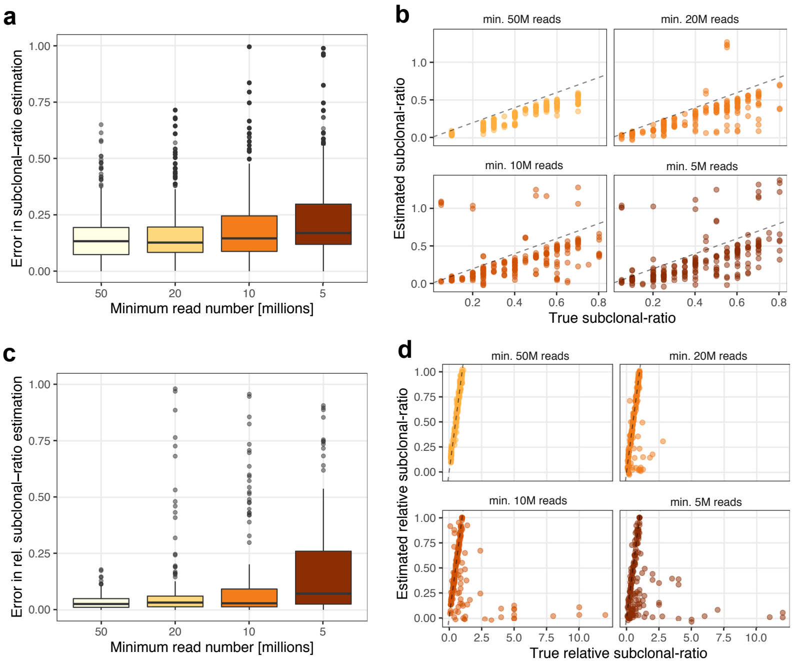


Figure S6: Subclonal-ratio estimation of *in silico* cell line mixtures, with CN values of each sample corrected using the theoretical rather than estimated tumour purity value. **(a,c)** Error in absolute (a) and relative (c) subclonal-ratio estimation of 200 datasets with minimum required read count indicated on the x axis. Box-plots represent: center line, median; box limits, upper and lower quartiles; whiskers, 1.5x interquartile range; points, outliers. **(b,d)** True and estimated subclonal-ratio of the datasets in (a) and (c).

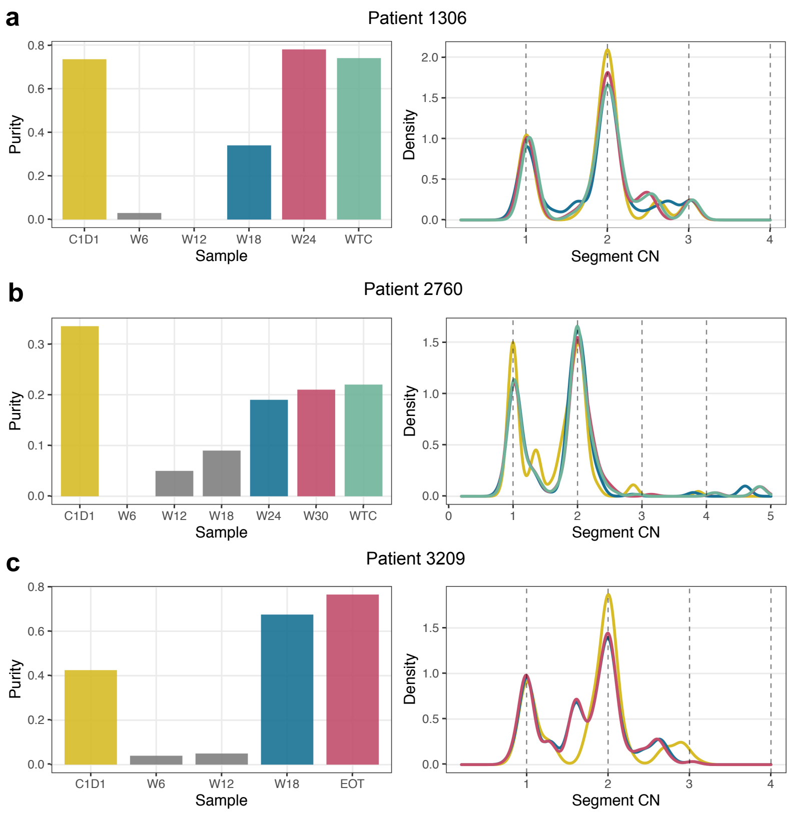


Figure S7: Tumour purity of cfDNA samples from lung cancer patients **(a)** 1306, **(b)** 2760, and **(c)** 3209. The left panel shows the purity estimate for each patient sample. Samples removed from downstream analysis are shown in grey. The right panel shows the purity-corrected CN distribution of non-discarded samples, coloured according to the bars on the left.
